# Rapid decay of host basal mRNAs during SARS-CoV-2 infection perturbs host antiviral mRNA biogenesis and export

**DOI:** 10.1101/2021.04.19.440452

**Authors:** James M. Burke, Laura A. St Clair, Rushika Perera, Roy Parker

**Affiliations:** Department of Biochemistry, University of Colorado Boulder, Boulder, Colorado, 80303; Howard Hughes Medical Institute, University of Colorado Boulder, Boulder, Colorado, 80303; BioFrontiers Institute, University of Colorado Boulder, Boulder, Colorado, 80303; Center for Vector-Borne and Infectious Diseases, Department of Microbiology, Immunology and Pathology, Colorado State University, Fort Collins, CO 80523, USA; Center for Metabolism of Infectious Diseases, Colorado State University, Fort Collins, CO 80523, USA

## Abstract

A key feature of the mammalian innate immune response to viral infection is the transcriptional induction of interferon (IFN) genes, which encode for secreted proteins that prime the antiviral response and limit viral replication and dissemination. A hallmark of severe COVID-19 disease caused by SARS-CoV-2 is the low presence of IFN proteins in patient serum despite elevated levels of *IFN*-encoding mRNAs, indicative of post-transcriptional inhibition of IFN protein production. Herein, we show SARS-CoV-2 infection limits type I and type III IFN biogenesis by preventing the release of mRNA from their sites of transcription and/or triggering their nuclear degradation. In addition, SARS-CoV-2 infection inhibits nuclear-cytoplasmic transport of *IFN* mRNAs as a consequence of widespread cytosolic mRNA degradation mediated by both activation of the host antiviral endoribonuclease, RNase L, and by the SARS-CoV-2 protein, Nsp1. These findings argue that inhibition of host and/or viral Nsp1-mediated mRNA decay, as well as IFN treatments, may reduce viral-associated pathogenesis by promoting the innate immune response.

## Introduction

The severe acute respiratory syndrome coronavirus 2 (SARS-CoV-2) is the cause of the COVID-19 pandemic. Since SARS-CoV-2 will remain endemic in human populations (Lavine et al., 2021), development of COVID-19 treatments is paramount. Several clinical trials are currently underway that modulate the innate immune response to treat COVID-19, including treatment with interferon (IFN) proteins (NCT04350671; NCT04388709; CT04647695; NCT04552379). However, the innate immune response to SARS-CoV-2 infection is not well-understood.

During the innate immune response to viral infection, the detection of viral double-stranded RNA (dsRNA) by host cells results in the transcriptional induction of mRNAs encoding for cytokines, including type I and type III IFNs, which are exported to the cytoplasm where they are translated (Jensen and Thomsen, 2012; Ivashkiv and Donlin, 2014; Lazear et al., 2019). These proteins are secreted from infected cells and prime an antiviral state in both infected and non-infected cells via autocrine and paracrine signaling. This limits viral replication capacity and promotes the function of innate and adaptive immune cells at sites of infection, which reduces viral loads and limits viral dissemination to secondary sites of infection.

Despite the potent antiviral activities of IFNs, it is currently controversial whether IFNs promote COVID-19 disease via their pro-inflammatory functions, or whether the low production of IFNs in response to SARS-CoV-2 contributes to COVID-19 disease progression. In support of the former, *IFN*-encoding mRNAs are elevated in patients with severe COVID-19 symptoms (Lee et al., 2020; Wilk et al., 2020; Zhou et al., 2020). In support of the latter, IFN proteins are relatively low in patients with severe COVID-19 symptoms (Blanco-Melo, et al., 2020; Hadjadj et al., 2020). While seemingly contradictory, these findings are nonetheless consistent with observations that SARS-CoV-2 induces transcription of IFNs, but antagonizes IFN protein production (Lei et al., 2020; Li et al., 2020). How IFNs are post-transcriptionally inhibited during SARS-CoV-2 infection is unknown.

Recently, we demonstrated that activation of the host antiviral endoribonuclease RNase L results in widespread degradation of mRNA (Burke et al., 2019), which in turn inhibits the nuclear export of *IFN* mRNAs, limiting IFN protein production (Burke et al., 2021). Herein, we analyze SARS-CoV-2 replication and its effect on host cells at the single-cell and single-molecule level. We show that SARS-CoV-2 infection similarly leads to rapid decay of host basal mRNAs triggering the inhibition of the nuclear export of *IFN* mRNAs. Unexpectedly, these phenomena can occur independently of RNase L activation via mRNA degradation mediated by the SARS-CoV-2-encoded Nsp1 protein. We also observe that SARS-CoV-2 infection limits the biogenesis of IFN mRNAs by reducing their release from their sites of transcription and/or triggering their nuclear degradation. These findings have implications for transcriptomic analyses of SARS-CoV-2 infection, current IFN drug trials, and the development of drugs to inhibit SARS-CoV-2-Nsp1-mediated mRNA decay.

## Results

### Generation of single-molecule SARS-CoV-2 mRNA visualization

To test the hypothesis that RNase L-mediated inhibition of mRNA export inhibits IFN protein production in response to SARS-CoV-2, we first transduced parental (WT) and RNase L knockout (RL-KO) A549 lung carcinoma cell lines with an ACE2-encoding lentivirus to make them permissive to SARS-CoV-2 infection (Burke et al., 2019) (Fig. S1A). We confirmed several RNase L-dependent phenotypes in response to poly(I:C) lipofection in WT^ACE2^ but not RL-KO^ACE2^ cells (Burke et al., 2019; Burke et al., 2020, Burke et al., 2021), including degradation of *GAPDH* mRNA, the generation of small stress granule-like foci (RLBs), inhibition of stress granule assembly, PABP translocation to the nucleus, and nuclear retention of *IFNB* mRNA (Fig. S1B,C). This demonstrates that these A549 cells expressing the ACE2 receptor have a normal innate immune response to dsRNA.

To identify SARS-CoV-2-infected cells, we generated single-molecule in situ hybridization (smFISH) probe sets that target the ORF1a, ORF1b, or N regions of the SARS-CoV-2 genomic mRNA (Fig. 1A). The ORF1a and ORF1b probes would be expected to detect the full-length (FL) genome, whereas the N probes would detect both the FL-genome and sub-genomic (SG) mRNAs (Fig. 1A).

**Fig. 1.**
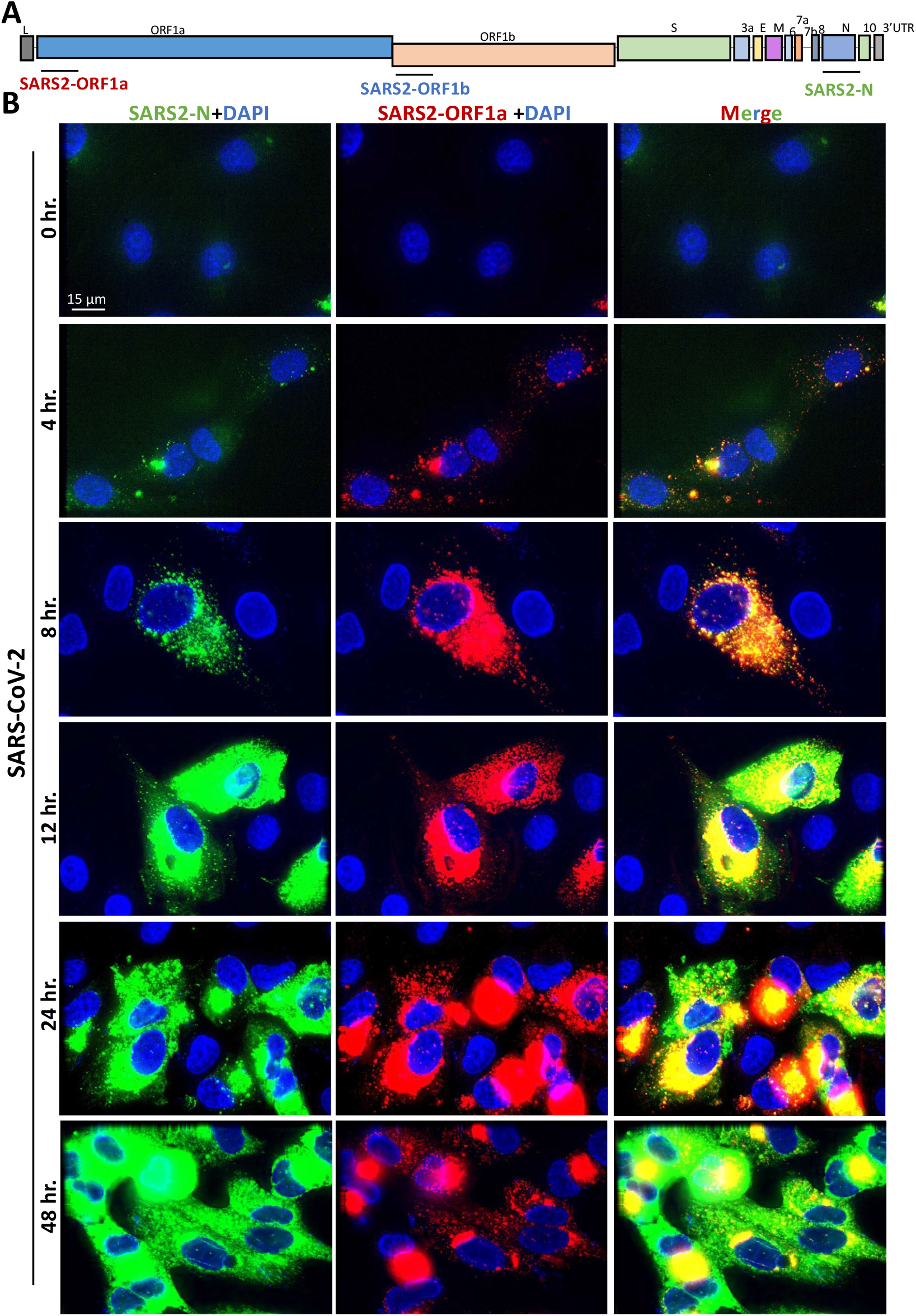
Single-molecule analysis of SARS-CoV-2 genomic and sub-genomic RNAs. (A) Schematic to show the location of smFISH probe sets targeting the different regions of SARS-CoV-2 mRNA. The ORF1a and ORF1b target the full-length genome, whereas the S and N probe sets target both the full-length genome and various sub-genomic RNAs. (B) smFISH for SARS-CoV-2 full length genome (ORF1a probes) and sub-genomic RNAs (N probes) at indicated times post-infection with SARS-CoV-2 (MOI=5).

We co-stained A549-WT^ACE2^ cells at multiple times post-infection with ORF1a and N smFISH probes. By four hours post-infection, we observed small and dispersed foci (~100 copies/cell) that co-stained for ORF1a and N RNA, which we suggest are individual SARS-CoV-2 genomes/full-length mRNAs (Fig. 1B and Fig. S2A). In addition, we observed larger structures that contain multiple genomes, which are likely replication factories (RFs) and/or concentrated sites of translation or mRNA processing. At eight hours post-infection, SARS-CoV-2 genome copies increased ~10-fold (to ~1000 copies/cell) and sub-genomic RNAs became abundant throughout the cell (Fig. 1B and Fig. S2B). From twelve through forty-eight hours post-infection, large RFs concentrated with full-length genome (fluorescent intensity was generally saturating) localized to the perinuclear region of the cell (Fig. 1B). At these later times, sub-genomic RNAs (N probes) were more abundant, as these N-positive RNAs only partially localized to the RFs and were mostly dispersed throughout the cytoplasm (Fig. 1B and Fig. S2B).

### SARS-CoV-2 infection triggers rapid degradation of host basal mRNAs independently of RNase L

To examine if SARS-CoV-2 infection activated RNase L-mediated mRNA decay, we stained for host *GAPDH* and *ACTB* basal mRNAs by smFISH. We observed a substantial reduction in *GAPDH* and *ACTB* mRNAs in SARS-CoV-2-infected cells WT^ACE2^ cells as early as eight hours post-infection (Fig. 2A,B and Fig. S2). Unexpectedly, we also observed a reduction in *GAPDH* and *ACTB* mRNAs in SARS-CoV-2-infected RL-KO^ACE2^ cells (Fig. 2C,D), indicating that the reduction in host basal mRNAs in response to SARS-CoV-2 infection can occur independently of RNase L.

**Fig 2.**
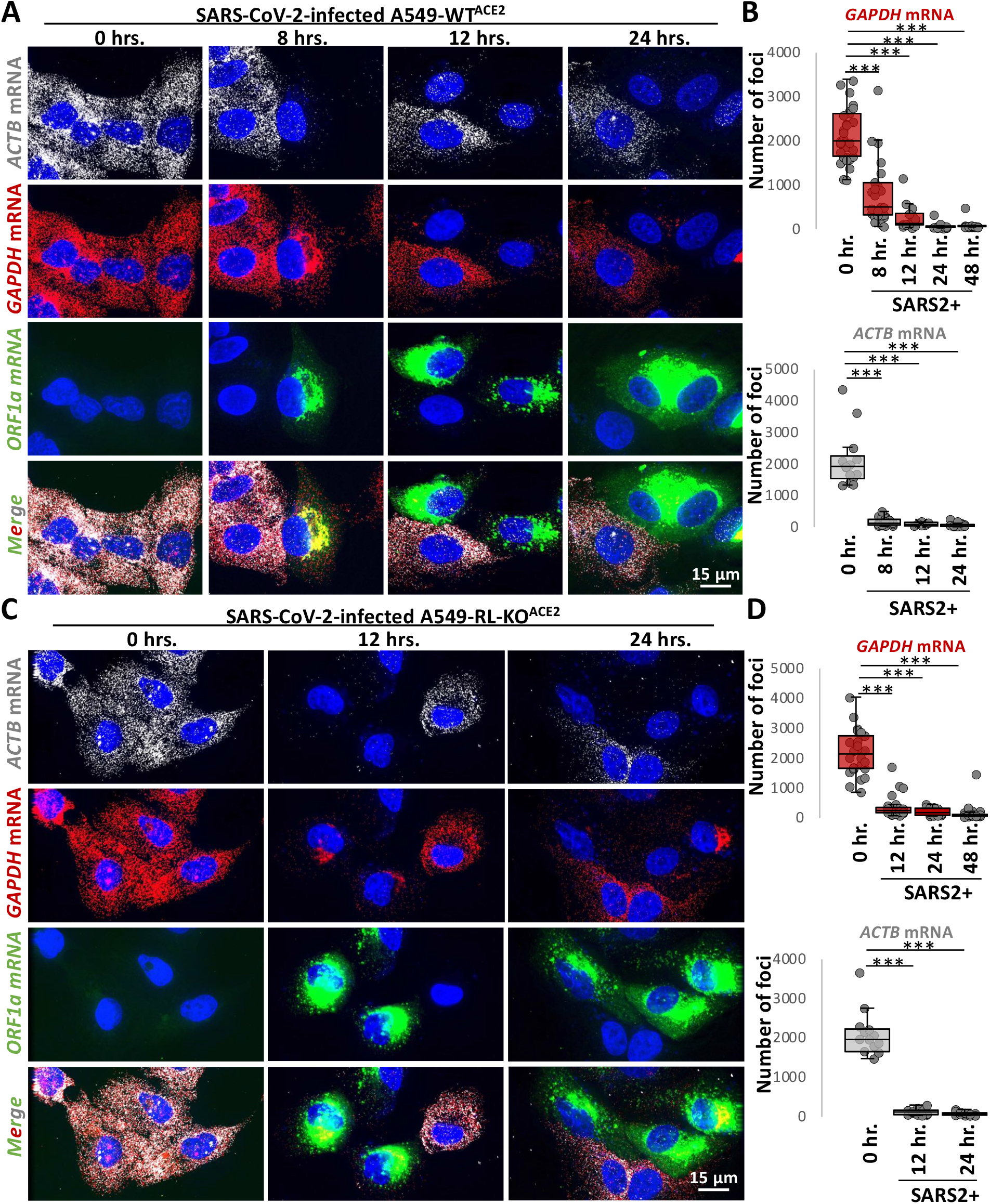
Host RNAs are rapidly degraded in response to SARS-CoV-2 infection, independently of RNase L. (A) smFISH for host *GAPDH* and *ACTB* mRNAs and SARS-CoV-2 full-length genome (ORF1b) at indicated times post-infection with SARS-CoV-2 (MOI=5) in WT^ACE2^ A549 cells. (B) Graphs show quantification of *GAPDH* and *ACTB* mRNAs as represented in above images. (C and D) Similar to (A and B) but in RL-KO^ACE2^ A549 cells.

However, several observations indicate that RNase L is activated by SARS-CoV-2 infection. First and consistent with RNase L reducing SARS-CoV-2 replication by ~4-fold (Li et al., 2020), we observed that RNase L reduced both FL-genome and N-RNA by ~3-fold as compared to the RL-KO^ACE2^ cells (Fig. S3A-C). Second, we observed robust RNase L-dependent accumulation of PABP in the nucleus by twenty-four hours post-infection (Fig. S3D), which is a previously reported consequence of RNase L activation (Burke et al., 2019). In contrast, despite widespread mRNA degradation in the RL-KO^ACE2^ cells, PABP did not translocate to the nucleus (Figure S3D). Lastly, we observed small punctate PABP-positive foci consistent with RLBs (<u>R</u>Nase <u>L</u>-dependent <u>b</u>odies) (Burke et al., 2019; Burke et al., 2020) (Fig. S3E).

Combined, these data indicate that SARS-CoV-2 infection leads to widespread degradation of host mRNAs both by the activation of RNase L, and by a second RNase L-independent mechanism.

### SARS-CoV-2 Nsp1 expression is sufficient for degradation of host basal mRNAs

The degradation of host basal mRNAs in the RL-KO^ACE2^ cells could either be an RNase L-independent host response or mediated by viral proteins. We hypothesized that either the Nsp1 or Nsp15 proteins encoded by SARS-CoV-2 could be responsible for host mRNA decay since Nsp1 can reduce host mRNA levels during coronavirus infection, possibly by inhibiting their translation (Narayanan et al., 2008; Schubert et al., 2020), and Nsp15 is an endoribonuclease that processes viral RNA (Bhardwaj et al., 2008), but could potentially cleave host mRNAs.

We generated flag-tagged SARS-CoV-2 Nsp1 or Nsp15 expression vectors and transfected them into U2-OS cells (Fig. 3A,B). In cells transfected with flag-Nsp1 (identified by staining for the flag epitope), both *ACTB* and *GADPH* mRNAs were strongly reduced in comparison to cells that did not stain for flag or cells transfected with empty vector (Fig. 3C,D). Expression of flag-Nsp15 did not result in notable reduction of *ACTB* and *GADPH* mRNAs (Fig. 3C,D). These data indicate that expression of SARS-CoV-2 Nsp1 protein is sufficient to initiate the widespread degradation of host basal mRNAs, arguing that Nsp1 contributes to host mRNA degradation during SARS-CoV-2 infection.

**Figure 3.**
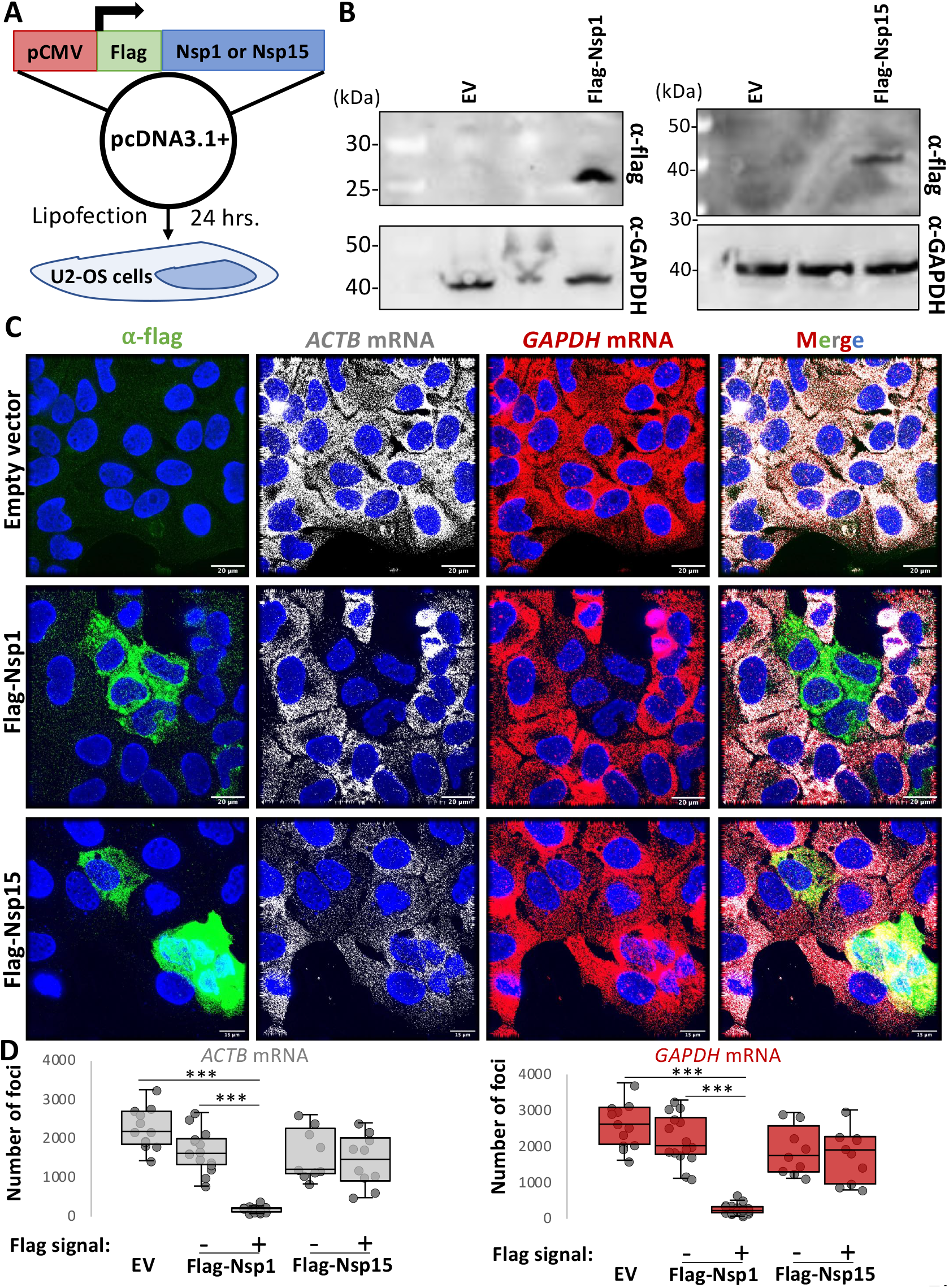
SARS-CoV-2 Nsp1 expression is sufficient for degradation of host basal mRNAs. (A) Schematic of flag-tagged SARS-CoV-2 protein expression vector transfected into U2-OS cells. (B) Immunoblot for flag confirmed expression of flag-tagged Nsp1 and Nsp15 expression at expected size (Nsp1 ~20 kDa; Nsp15 ~40kDa) in cells transfected with respective expression vectors but not empty vector (EV). (C) Immunofluorescence assay for flag and smFISH for *ACTB* and *GAPDH* mRNAs in U2-OS twenty-four hours post transfection with either pcDNA3.1+ (empty vector; EV), flag-Nsp1, or flag-Nsp15 expression vectors. (D) Quantification of *ACTB* and *GAPDH* mRNAs as represented in (C). Statistical significance (*p<0.05; **p<0.005; ***p<0.0005) was determined by t-test.

### Alterations to type I and type III *IFN* mRNA biogenesis during SARS-CoV-2 infection

The observations that SARS-CoV-2 both activates RNase L (Fig. S3) and also promotes decay of host basal mRNAs via Nsp1 (Fig. 2 and 3) suggests that *IFN* mRNAs might be retained in the nucleus due to an mRNA export block triggered by widespread cytosolic RNA degradation (Burke et al., 2021). Given this possibility, we examined the expression of *IFN* mRNAs by smFISH during SARS-CoV-2 infection. These experiments revealed three important insights into how SARS-CoV-2 affects IFN production.

#### SARS-CoV-2 infection induces RLR-MAVS-IRF3/7 signaling similar to dsRNA mimics

We observed that SARS-CoV-2 infection generally triggers RLR-MAVS-IRF3/7 signaling, leading to transcriptional induction of *IFN* genes. This is based on the observation that 45% of SARS-CoV-2-infected A549-WT^ACE2^ cells stain positive for abundant disseminated *IFNB1* mRNA and/or nascent *IFNB1* transcripts at *IFNB1* genomic loci, referred to as transcriptional foci (Burke et al., 2019) (Fig. 4A,C). The lack of *IFNB1* induction in 55% of SARS-CoV-2-infected cells is likely due to the inherent heterogeneity of the innate immune response in A549 cells (Burke et al., 2019), since a similar, albeit slightly lower, percentage (37%) of A549-WT cells that were transfected with poly(I:C) (as determined by RNase L activation) did not induce *IFNB1* expression (Fig. 4B,C). These data indicate that SARS-CoV-2 infection often leads to RLR-MAVS-IRF3/7-mediated *IFN* gene induction in A549 cells.

**Fig. 4.**
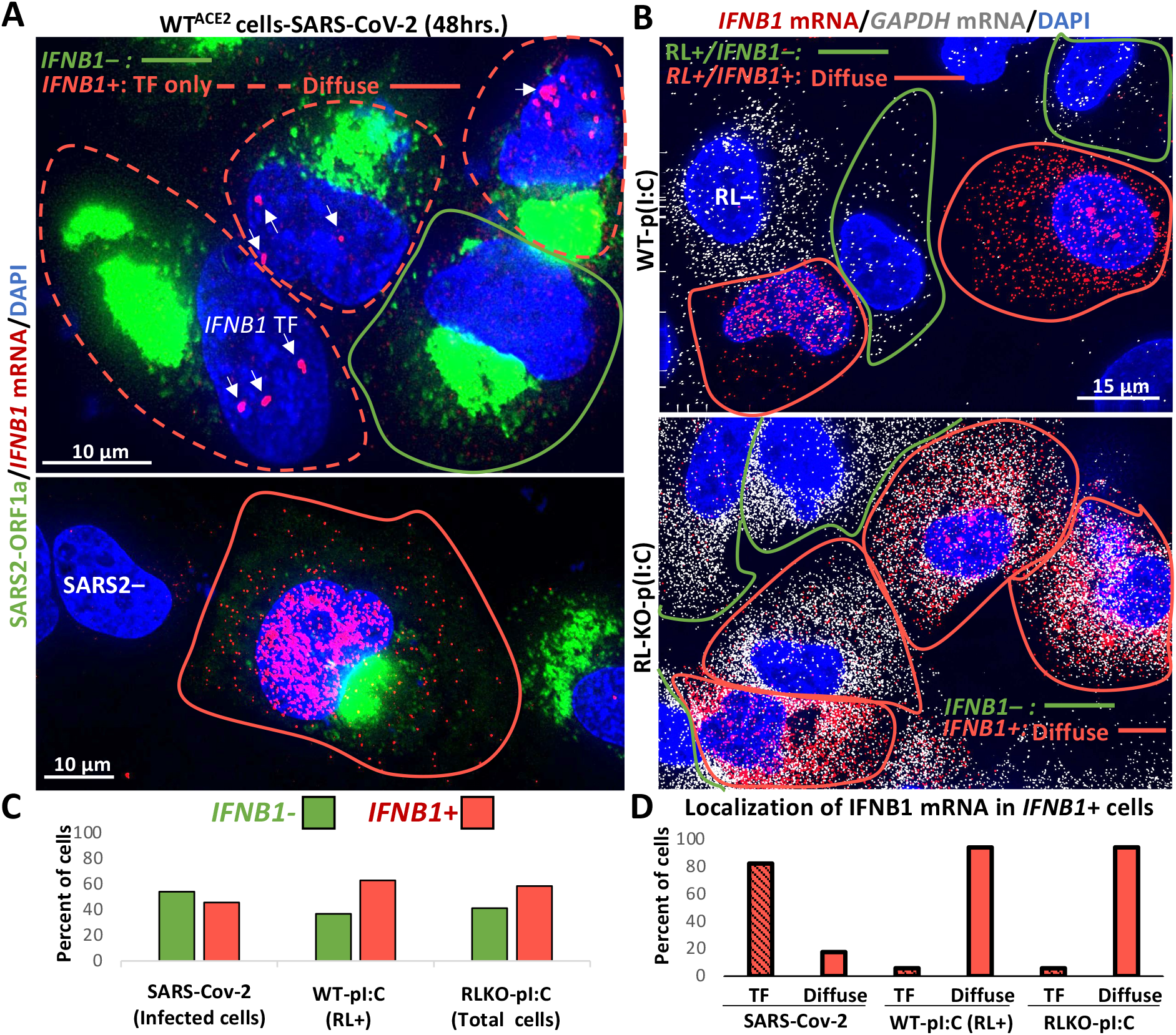
*IFN* mRNAs are retained at the site of transcription during SARS-CoV-2 infection. (A) smFISH for *IFNB1* mRNA and SARS-CoV-2 ORF1a forty-eight hours post-infection. Two fields of view are shown. In the top image, 45% of SARS-CoV-2-positive cells stain for *IFNB1* (cell boundary marked by red line), whereas 55% do not (cell boundary marked by green line). Eighty-two percent of cells that contain *IFNB1* transcriptional foci (TF) lack abundant disseminated *IFNB1* mRNA (dashed red line). The lower image shows a SARS-CoV-2-infected cell that contains abundant and diffuse *IFNB1* mRNA in the nucleus and cytoplasm (solid red line), which constitute less than 18% of cells that induce *IFNB1* (contain IFNB1 transcriptional foci or mRNA). Cells that do not stain for SARS-CoV-2 are labeled SARS2-negative (SARS2-). (B) smFISH for *IFNB1* mRNA and *GAPDH* mRNA sixteen hours post-poly(I:C) transfection in WT and RL-KO A549 cells (images of individual staining shown in Fig. S4B). In WT cells, 12% do not activate RNase L (RL-). Of the 88% of cells that activate RNase L, 63% (55% of total cells) also induce abundant and disseminated *IFNB1* mRNA (cell boundary marked by red line), whereas 37% of RL+ cells do not induce IFNB1 (cell boundary marked by green line). Fifty-nine percent of RL-KO cells induce abundant disseminated IFNB1 mRNA (cell boundary marked by red line), whereas 41% do not (cell boundary marked by green line). (C) Histograms quantifying the percent of SARS-CoV-2 infected cells, poly(I:C)-transfected WT cells that activate RNase L (GAPDH mRNA-negative cells), and poly(I:C)-transfected RL-KO cells that induce *IFNB1*, as represented in (A and B). (D) Histograms quantifying the percent of *IFNB1*-positive cells in which IFNB1 smFISH staining is predominantly localized to *IFNB1* transcriptional foci (TF) or diffuse.

#### SARS-CoV-2 infection leads to retention of IFN mRNAs at sites of transcription and/or nuclear degradation

Importantly, several observations suggest that SARS-CoV-2 either disrupts *IFNB1* mRNA release from sites of transcription and/or causes nuclear degradation of *IFNB1* mRNA (Fig 4A-D). Specifically, of the SARS-CoV-2-infected cells that induced *IFNB1*, greater than 82% contained *IFNB1* transcriptional foci but lacked abundant diffuse *IFNB1* mRNAs (Fig. 4A,D). In these cells, *IFNB1* mRNAs were few in number and limited to the vicinity of the *IFNB1* transcriptional foci (Fig. 4A). Less than 18% of SARS-CoV-2 infected cells displayed abundant *IFNB1* mRNAs that had disseminated away from the *IFNB1* site of transcription (Fig. 4A,D). We observed a similar effect staining for *IFNL1* mRNA (Fig. S4A).

Our data indicate that the inability of *IFNB1* mRNA to disseminate away from *IFNB1* transcriptional foci during SARS-CoV-2 infection is not typical of *IFNB1* induction nor a consequence of RNase L activation. Specifically, of the WT or RL-KO A549 cells that induce *IFNB1* in response to poly(I:C) lipofection (Fig. 4B,C), greater than 94% displayed widespread dissemination of *IFNB1* mRNA in the nucleus and/or in the cytoplasm, with very few cells (<6%) displaying clearly identifiable *IFNB1* transcriptional foci but lacking disseminated *IFNB1* mRNA (Fig. 4B,D and S4B). Since most WT cells activated RNase L in response to poly(I:C), and all WT cells that induced *IFNB1* also activated RNase L (Fig. 4B,C), RNase L activation does not cause retention of *IFNB1* mRNA at *IFNB1* transcriptional foci. Further supporting that RNase L does not cause this effect, we observed this phenomenon in SARS-CoV-2-infected RL-KO^ACE2^ cells (Fig. S4C).

These data argue that SARS-CoV-2 infection, as opposed to *IFNB1* induction or RNase L activation, mediates the inhibition of *IFNB1* mRNA release from the site of transcription. One possible mechanism for this effect is an alteration in RNA processing, which can prevent the release of mRNAs from transcriptional foci (Hilleren et al., 2001). Given this, we examined whether *IFNB1* mRNAs were similarly retained at transcriptional foci during influenza A virus (IAV) infection, which is known to perturb host mRNA processing and export (Fortes et al., 1994; Hayman et al., 2006; Nemeroff et al., 1998). Indeed, we observed a similar effect during IAV infection, whereby of the 32% of IAV-infected cells that induced *IFNB1* (presence of *IFNB1* foci), the majority (96%) lacked abundant disseminated *IFNB1* mRNA but contained intense *IFNB1* transcriptional foci (Fig. S4C). Therefore, the lack of disseminated *IFNB1* mRNAs in response to SARS-CoV-2 infection indicates that SARS-CoV-2, as wells as IAV, leads to a viral-mediated block in release of *IFN* mRNAs from their sites of transcription and/or rapid degradation of *IFN* mRNAs upon leaving the site of transcription.

#### SARS-CoV-2 infection blocks nuclear export of IFN mRNAs

A second mechanism we observed by which SARS-CoV-2 infection limits IFN protein production is by a block in the transport of *IFN* mRNAs from the nucleus to the cytoplasm. The critical observation is that in SARS-CoV-2 infected cells that induced *IFNB1* and displayed abundant, diffuse *IFNB1* mRNA, 75% retained the majority (>65%) *IFNB1* mRNAs in the nucleus (Fig. 5A,B). Similar results were seen with IFNL1 mRNA (Fig. S4E). The nuclear retention of *IFN* mRNAs during SARS-CoV-2 infection is similar to that observed in response to RNase L activation during poly(I:C) lipofection or dengue virus serotype 2 (DENV2) infection (Burke et al., 2021).

**Fig. 5.**
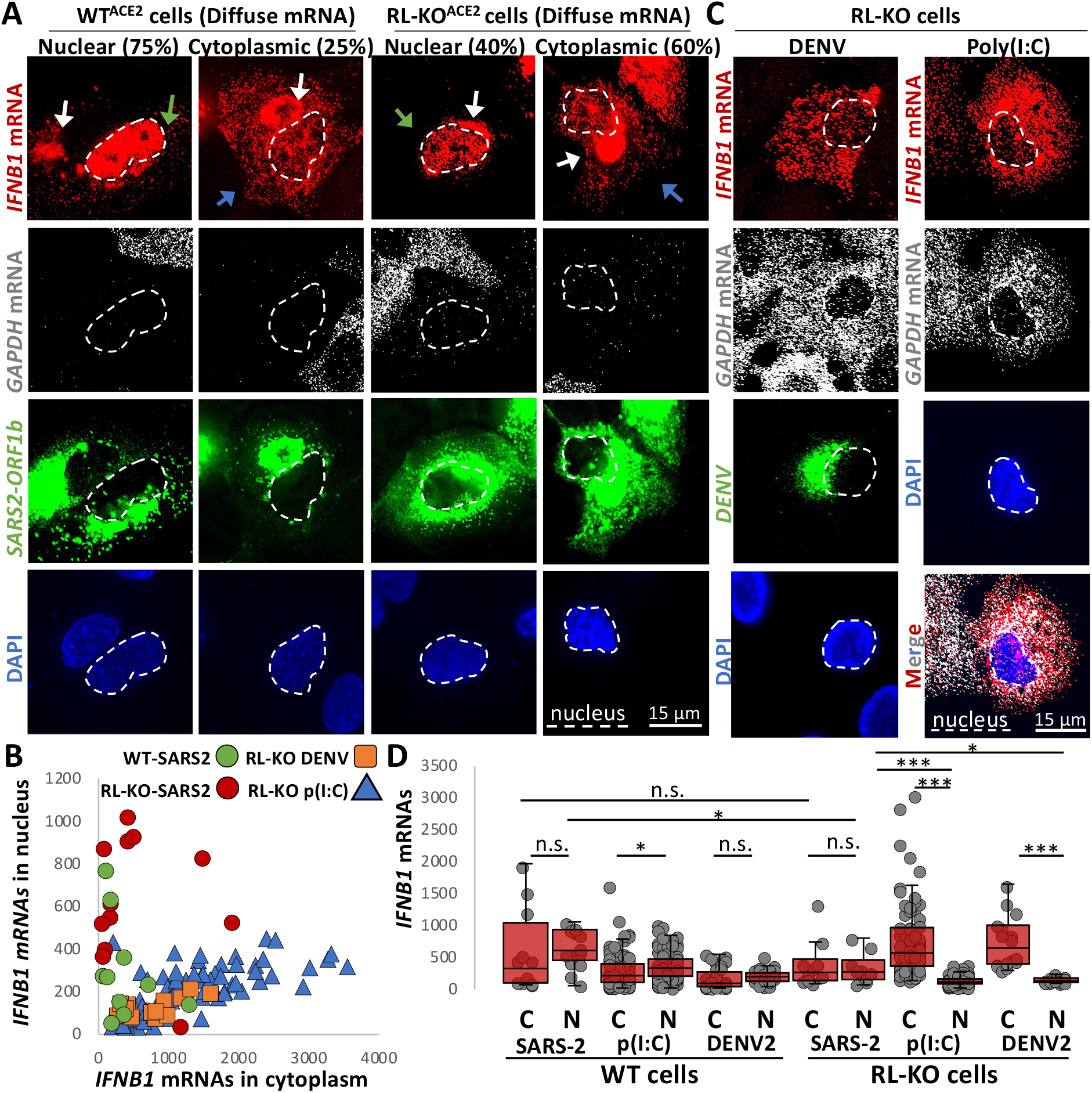
Nuclear-cytoplasmic transport of *IFN* mRNAs is inhibited during SARS-CoV-2 infection. (A) smFISH for *IFNB1* mRNA, *GAPDH* mRNA, and SARS-CoV-2 *ORF1b* mRNA in WT^ACE2^ and RL-KO^ACE2^ cells forty-eight hours post-infection with SARS-CoV-2 (MOI=5). Spectral crossover from the SARS-CoV-2 ORF1b RF into the IFNB mRNA channel is indicated be white arrows. The green arrows indicate cells in which *IFNB* mRNA is retained in the nucleus. The blue arrows indicate cells in which *IFNB* mRNA is localized to the cytoplasm. (B) Scatter plot quantifying IFNB1 mRNA in the nucleus (y-axis) and in the cytoplasm (x-axis) in individual WT^ACE2^ or RL-KO^ACE2^ cells infected with SARS-CoV-2, or RL-KO cells 48 hrs. post-infection with DENV or 8 hours post-transfection with poly(I:C). (C) Representative smFISH for *IFNB1* and *GAPDH* mRNAs in RL-KO A549 cells forty-eight hours post infection with DENV (MOI=0.1) or 16 hours post-transfection with poly(I:C). (D) Quantification of *IFNB1* mRNA via smFISH in the nucleus (N) or cytoplasm (C) of either WT^ACE2^ or RL-KO^ACE2^ cells infected with SARS-CoV-2, and WT or RL-KO cells transfected with poly(I:C) or infected with DENV2 as represented in (A and C). Poly(I:C) and DENV2 data was obtained from Burke et al., 2021. Statistical significance (*p<0.05; **p<0.005; ***p<0.0005) was determined by t-test.

However, nuclear retention of *IFN* mRNAs during SARS-CoV-2 infection can occur independently of RNase L activation. The key observation is that we observed nuclear retention of *IFNB1* mRNA in SARS-CoV-2-infected RL-KO^ACE2^ cells (Fig. 5A,B). This is in contrast to poly(I:C) lipofection or DENV2 infection in RL-KO cells, in which *IFNB1* mRNA is predominantly localized to the cytoplasm (Fig. 5B,C) (Burke et al., 2021).

A notable difference between SARS-CoV-2 infection and either poly(I:C) lipofection or DENV infection in RL-KO cells is that basal mRNAs are only degraded in the context of SARS-CoV-2 infection (Fig. 5A,C). This suggests that SARS-CoV-2-mediated mRNA decay might be responsible for inhibiting the nuclear export of *IFNB1* mRNAs, similar to RNase L-mediated mRNA decay (Burke et al., 2021). To better assess this model, we compared nuclear and cytoplasmic *IFNB1* mRNA levels during SARS-CoV-2 infection, poly(I:C) lipofection, or DENV2 infection in both WT and RL-KO cells.

This analysis supports that SARS-CoV-2-mediated mRNA decay, similar to RNase L-mediated mRNA decay, inhibits IFNB1 mRNA export. Specifically, while median *IFNB1* mRNA levels in the cytoplasm are ~8-fold higher than in the nucleus of RL-KO cells infected with DENV2 or transfected with poly(I:C), they are equivalent in SARS-CoV-2-infected RL-KO cells (Fig. 5D). Moreover, the ratio of nuclear to cytoplasmic *IFNB1* mRNA levels in SARS-CoV-2-infected RL-KO cells is comparable to WT cells infected with SARS-CoV-2, DENV2, or transfected with poly(I:C) (Fig. 5D).Thus, the high percentage of cells displaying nuclear retention of *IFNB1* mRNA is specific to scenarios in which widespread decay of host mRNA occurs, including RNase L activation (Burke et al., 2021) and SARS-CoV-2 infection (Fig. 2).

Interestingly, in a fraction of SARS-CoV-2-infected WT^ACE2^ or RL-KO^ACE2^ cells (25% and 40% respectively) displaying diffuse *IFNB1* mRNA staining, *IFNB1* mRNA was abundant in the cytoplasm despite robust decay of *GAPDH* mRNA (Fig. 5A,B). These data indicate that the *IFNB1* mRNA at least partially evades both RNase L- and Nsp1-mediated mRNA decay mechanisms during SARS-CoV-2 infection when the *IFNB1* mRNA is successfully exported to the cytoplasm. This is similar to results seen with activation of RNase L either by poly(I:C) transfection or DENV2 infection (Burke et al., 2019; Burke et al., 2021).

## Discussion

Several observations support that SARS-CoV-2 infection perturbs *IFN* mRNA biogenesis, limiting *IFN* mRNAs from reaching the cytoplasm where they can be translated (Fig. 6). First, *IFN* genes are induced in ~45% of SARS-CoV-2-infected cells, indicating that RLR-MAVS-IRF3/7 signaling is initiated by SARS-CoV-2, consistent with a previous report (Li et al., 2020). However, we observed that both type I- and type III IFN-encoding mRNAs predominately localize to their transcriptional sites in a majority of cells (Fig. 4A,D and S4A). Since this effect is atypical of *IFN* induction and not a consequence of RNase L activation in response to poly(I:C) (Fig. 4B,D), we suggest that SARS-CoV-2 inhibits some aspect of RNA processing or an early step of mRNA export, either of which is necessary for efficient release of stable mRNAs from transcription sites (Hilleren et al., 2001). Notably, we observed a similar phenomenon occurring during IAV infection (Fig. S4D), which is known to perturb host mRNA processing and export (Fortes et al., 1994; Hayman et al., 2006; Nemeroff et al., 1998).

**Fig. 6.**
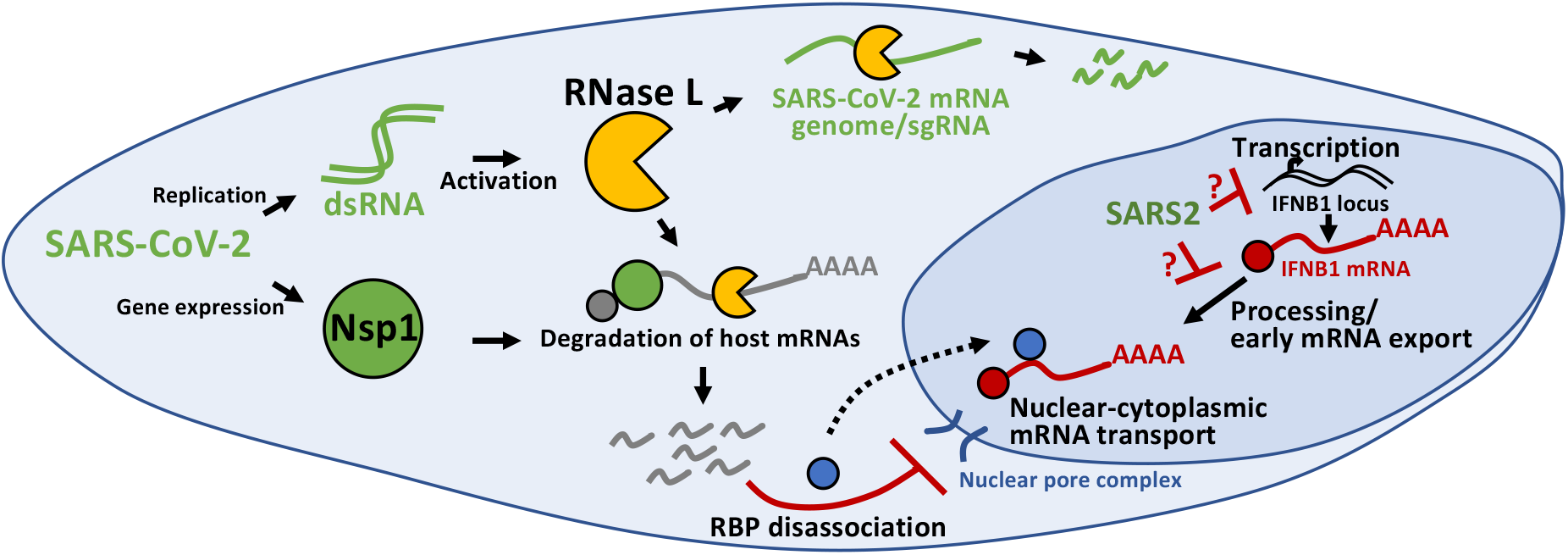
Inhibition of antiviral mRNA biogenesis during SARS-CoV-2 infection. Schematic modeling how antiviral mRNA biogenesis is inhibited during SARS-CoV-2 infection. SARS-CoV-2 replication generates double-stranded RNA (dsRNA), which leads to RNase L activation. RNase L-mediated mRNA decay reduces SARS-CoV-2 full-length mRNA genome and sub-genomic mRNAs. In addition, SARS-CoV-2 expresses the viral Nsp1 protein. Both RNase L activation and Nsp1 expression result in rapid and widespread decay of host basal mRNAs. While RNase L directly cleaves mRNAs, the mechanism of Nsp1-mediated mRNA decay is unclear. The degradation of host mRNAs results in release of RNA-binding proteins (RBPs), and this perturbs late stages of nuclear-cytoplasmic RNA transport. The sequestration of antiviral mRNAs, such as *IFNB1* mRNA, in the nucleus prevents their association with ribosomes in the cytoplasm, reducing their translation for protein production. In addition, SARS-CoV-2 inhibits the transcription, an aspect of mRNA processing, or association with early mRNA export factors, and/or rapidly degrades dsRNA-induced antiviral mRNAs, such as *IFNB1* mRNA. The result of this is the inability of *IFNB1* mRNAs to exit the site of *IFNB1* transcription, preventing their transport to the cytoplasm and reducing their translation.

In addition to inhibition of early mRNA processing/export, we observed phenotypes consistent with the inhibition of the late steps of nuclear-cytoplasmic mRNA transport during SARS-CoV-2 infection. Specifically, we observed that the majority of cells in which IFN mRNAs were released from the sites of transcription retained those *IFN* mRNAs within, but disseminated throughout, the nucleus in SARS-CoV-2-infected cells (Fig. 5A). However, the mRNA export block of *IFN* mRNAs is distinct from the accumulation of *IFN* mRNAs at transcriptional foci since a similar mRNA export block is triggered by RNase L without reduction of transcription nor trapping of mRNAs at the transcription site in both poly(I:C)-treated and DENV2-infected cells (Burke et al., 2021) (Fig. 4A-D).

The inhibition of nuclear mRNA export by SARS-CoV-2 infection can be understood as a direct consequence of widespread mRNA degradation in the cytosol. The key observation is that we observed rapid and widespread decay of host basal mRNAs in response to SARS-CoV-2 (Fig. 2), which could be mediated by RNase L activation (Fig. S3) and/or the SARS-CoV-2 Nsp1 protein (Fig. 3). Moreover, since we have recently shown that RNase L-mediated mRNA decay inhibits mRNA export of *IFN* mRNAs (Burke et al., 2021), these data argue that either RNase L- or SARS-CoV-2-Nsp1-mediated mRNA decay leads to inhibition of host mRNA export. It should be noted that RNase L per se is not required for this export block since we observed *IFN* mRNAs trapped in the nucleus in RNase L knockout cells where widespread mRNA degradation is driven by Nsp1 (Figs. 3 and 5A,B,D). Regardless of the nuclease responsible for mRNA destruction, the nuclear retention of *IFN* mRNAs away from ribosomes would consequently reduce IFN protein production in response to SARS-CoV-2 infection.

Although the detailed mechanism of the mRNA export block is unknown, it appears to be a general consequence of any widespread cytosolic mRNA degradation. This mechanism is suggested by the observations that mRNA export blocks occur due to activation of RNase L (Burke et al., 2021), the nuclease SOX2 produced by Kaposi’s sarcoma-associated herpesvirus (KSHV) (Gilbertson et al., 2018; Glaunsinger et al., 2005; Kumar and Glaunsinger, 2010), and by degradation of mRNAs by Nsp1 in RL-KO cells (Figs. 3 and 4E). A likely explanation for the export block is that widespread cytosolic mRNA degradation leads to re-localization of numerous RNA binding proteins to the nucleus (Khong and Parker 2020, RNA; Burke et al., 2019; Kumar and Glaunsinger, 2010), which would then compete for the binding of export factors to mRNAs. Consistent with that hypothesis, overexpression of the mRNA export factor NXF1 (Nuclear RNA Export Factor 1) has been suggested to overcome an mRNA export block due to Nsp1 binding to NXF1 (Zhang et al., 2021). However, we anticipate that Nsp1 binding to NXF1 would not be required for inhibition of mRNA export in SARS-CoV-2 infected cells since anytime mRNAs are degraded via RNase L activation, which happens in SARS-CoV-2 infections (Fig. S3) (Li et al., 2020), there is a robust mRNA export block independent of any viral protein (Burke et al., 2021). Moreover, we observe inhibition of *IFNB1* mRNA export during SARS-CoV-2 infections, which is exported by CRM1-dependent export pathway and is independent of NXF1 (Fig. 5A) (Burke et al., 2021). An important issue for future research is to understand the factors that compete for mRNA export once cytosolic mRNAs are degraded.

Despite rapid degradation of host basal mRNAs, SARS-CoV-2 RNAs appeared to be largely unaffected since they increased over time and were only modestly reduced by RNase L (Figs. 1, S2, S3). Similarly, in cells in which *IFNB* mRNAs were exported to the cytoplasm, *IFNB1* mRNAs appeared to be stable since they were abundant despite complete decay of basal mRNAs (Fig. 5A), similar to *IFNB1* mRNA escaping RNase L-mediated mRNA decay (Burke et al., 2019). Importantly, this indicates that *IFN* mRNAs evade SARS-CoV-2-mediated mRNA decay mechanisms, making rescue of host mRNA processing and export a viable option for increasing IFN protein production.

## Materials and methods

### Cell culture

Parental and RNase L-KO (RL-KO) A549, U2-OS, and HEK293T cell lines are described in Burke et al., 2019. Cells were maintained at 5% CO_2_ and 37 degrees Celsius in Dulbecco’s modified eagle’ medium (DMEM) supplemented with fetal bovine serum (FBS; 10% v/v) and penicillin/streptomycin (1% v/v). Cells were routinely tested for mycoplasma contamination by the cell culture core facility. Cells were transfected with poly(I:C) HMW (InvivoGen: tlrl-pic) using 3-μl of lipofectamine 2000 (Thermo Fisher Scientific) per 1-ug or poly(I:C). African green monkey kidney cells (Vero E6, ATCC CRL-1586) were maintained at 5% CO_2_ and 37 degrees Celsius in DMEM supplemented with FBS (10% v/v), 2 mM non-essential amino acids, 2 mM l-glutamine, and 25 mM HEPES buffer.

### Plasmids

The flag-Nsp1 and flag-Nsp15 vectors were generated by ligating a g-block [Integrated DNA Technologies (IDT)] encoding for the flag and ORF of Nsp1 or Nsp15 between the *xho1* and *xba1* sites in pcDNA3.1+. Plasmids were sequence verified. The pLEX307-ACE2-puro plasmid was a gift from Alejandro Chavez and Sho Iketani (Addgene plasmid # 158448; http://n2t.net/addgene: 158448; RRID:Addgene_158448).

### Viral infections

SARS-CoV-2/WA/20/01 (GenBank MT020880) was acquired from BEI Resources (NR-52881) and used for all infections. The virus was passaged in Vero E6 cells, and viral titer was determined via plaque assay on Vero E6 as previously described in (Dulbecco et al., 1953). A multiplicity of infection (MOI) of 5 was used unless otherwise noted. All cell culture and plate preparation work were conducted under biosafety level 2 conditions, while all viral infections were conducted under biosafety level 3 conditions at Colorado State University. For infections, cells were seeded in 6-wells format onto cover slips. Twenty-four hours later, cell growth medium was removed, and cells were inoculated with SARS-CoV-2 at the indicated MOI for 1 hour at room temperature to allow for viral adherence. After incubation, viral inoculum was removed, cells were washed with 1X PBS, and DMEM supplemented with 2% FBS (v/v) was added to each well. Cells were fixed in 4% paraformaldehyde and phosphate-buffered solution (PBS) for 20 minutes, followed by three five-minute washes with 1X PBS, and stored in 75% ethanol. Following paraformaldehyde fixation, plates were removed from BSL3 facility, and stored at 4 degrees Celsius until staining. A549 cells were infected with dengue virus serotype 2 16681 strain at an MOI of 0.1, and with influenza A/Udorn/72 virus strain at MOI of 0.5, as described in Burke et al., 2021.

### Generation of ACE2-expressing cell lines

HEK293T cells (T-25 flask; 80% confluent) were co-transfected with 2.4-ug of pLenti-pLex307-ACE lentiviral transfer plasmid (Addgene: 158448), 0.8-ug of pVSV-G, 0.8-ug of pRSV-Rev, and 1.4-ug of pMDLg-pRRE using 20-ul of lipofectamine 2000. Media was collected at twenty-four- and forty-eight-hours post-transfection and filter-sterilized with a 0.45-um filter. To generate A549^ACE2^ line, cells were incubated for 1 hour with 1-ml of ACE2-encoding lentivirus with 10-ug of polybrene. Normal medium was then added to the flask and incubated for twenty-four hours. Medium was removed 24 hours post-transduction and replaced with selective growth medium containing 2-ug/ml of puromycin (Sigma-Aldrich). Selective medium was changed every three days. After one-week, selective medium was replaced with normal growth medium. Expression of ACE2 was confirmed via immunoblot analysis (protocol described in Burke et al., 2019) using Anti-Angiotensin Converting Enzyme 2 antibody [EPR4435(2) (Abcam: ab108252) at 1:1000.

### Single-molecule fluorescent in situ hybridization (smFISH) and immunofluorescence assays

smFISH was performed following manufacturer’s protocol (https://biosearchassets.blob.core.windows.net/assets/bti_custom_stellaris_immunofluorescence_seq_protocol.pdf) and as described in Burke et al., 2019 and Burke et al., 2021. GAPDH and ACTB smFISH probes labeled with Quasar 570 Dye (GAPDH: SMF-2026-1) or Quasar 670 Dye (GAPDH: SMF-2019-1) (ACTB: VSMF-2003-5) were purchased from Stellaris. Custom IFNB1, IFNL1, and SARS-CoV-2 smFISH probes were designed using Stellaris smFISH probe designer (Biosearch Technologies) available online at http://www.biosearchtech.com/stellaris-designer. Reverse complement DNA oligos were purchased from IDT (Extended data file 1). The probes were labeled with Atto-633 using ddUTP-Atto633 (Axxora: JBS-NU-1619-633), with ATTO-550 using 5-Propargylamino-ddUTP (Axxora; JBS-NU-1619-550), or ATTO-488 using 5-Propargylamino-ddUTP (Axxora; JBS-NU-1619-488) with terminal deoxynucleotidyl transferase (Thermo Fisher Scientific: EP0161) as described in (Gaspar et al., 2017).

For immunofluorescence detection, cells were incubated with Rabbit polyclonal anti-PABP antibody (Abcam: ab21060) and Mouse monoclonal anti-G3BP antibody (Abcam: ab56574) primary antibodies at 1:1000 for two hours, washed three times, and then incubated with Goat Anti-Rabbit IgG H&L (Alexa Fluor^®^ 647) (Abcam: ab150079) and Goat Anti-Mouse IgG H&L (FITC) (Abcam; ab97022) at 1:2000 for one hour. After three washes, cells were fixed and then smFISH protocol was performed.

### Microscopy and Image Analysis

Microscopy was performed as described in Burke et al., 2021. Briefly, cover slips were mounted on slides with VECTASHIELD Antifade Mounting Medium with DAPI (Vector Laboratories; H-1200). Images were obtained using a wide field DeltaVision Elite microscope with a 100X objective using a PCO Edge sCMOS camera. 10 Z planes at 0.2 um/section were taken for each image. Deconvoluted images were processed using ImageJ with FIJI plugin. Z-planes were stacked, and minimum and maximum display values were set in ImageJ for each channel to properly view fluorescence. Imaris Image Analysis Software (Bitplane) (University of Colorado-Boulder, BioFrontiers Advanced Light Microscopy Core) was used to quantify smFISH foci in nucleus and cytoplasm. Fluorescent intensity was quantified in ImageJ.

## Acknowledgments

The authors thank Dr. Carolyn Decker for valuable comments regarding the manuscript. Research reported in this publication was supported by the National Institute of Allergy and Infectious Diseases of the National Institutes of Health under Award Number F32AI145112 (J.M.B), funds from HHMI (Roy Parker), and support provided by the Office of the Vice President for Research and the Dept. of Microbiology, Immunology and Pathology at Colorado State University (Rushika Perera).

## Author contributions

J.M.B and Roy Parker conceived the project. L.A.S. performed SARS-CoV-2 infections. J.M.B. generated cell lines and plasmids, performed microscopy, and quantified microscopy data. J.M.B., L.A.S., Rushika Perera., and Roy Parker. interpreted data. J.M.B. and Roy Parker wrote the manuscript.

