## Supplemental Figures and Legends for "Rapid decay of host basal mRNAs during SARS-CoV-2 infection perturbs host antiviral mRNA biogenesis and export"

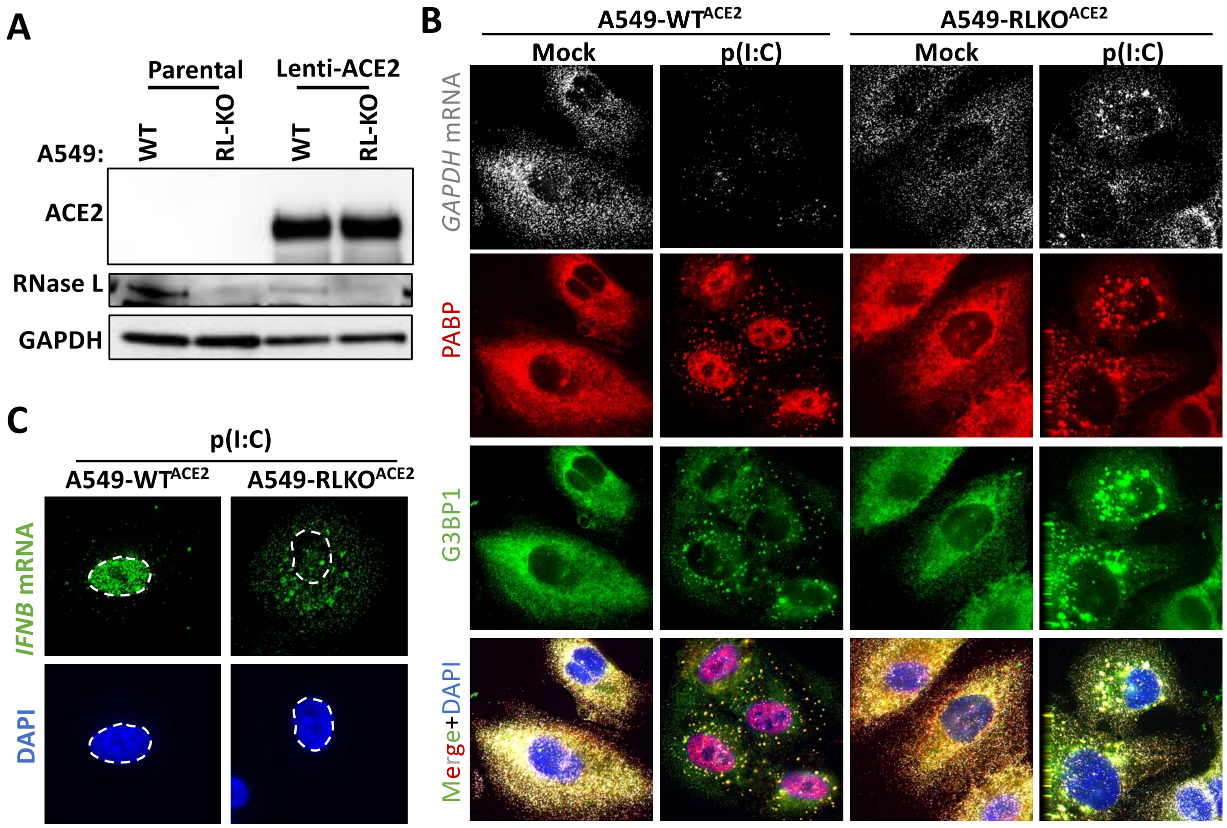


**Figure. S1. Generation and characterization of WT and RNase L-KO A549 cells that express ACE2.** (A) Immunoblot analysis to confirm ACE2 expression in parental (WT) and RNase L-KO (RL-KO) A549 cells. (B) Single-molecule fluorescent in situ hybridization (smFISH) for GAPDH mRNA and immunofluorescent assay for RNA-binding proteins PABP and G3BP1 that enrich in RNase L-dependent bodies (RLBs) in WT cells and stress granules in RL-KO cells four hours post-lipofection of poly(I:C). PABP accumulates in the nucleus in WT cells. (C) smFISH for *IFNB* mRNA in WT$^{ACE2}$ and RL-KO$^{ACE2}$ cells four hours post-lipofection of poly(I:C).

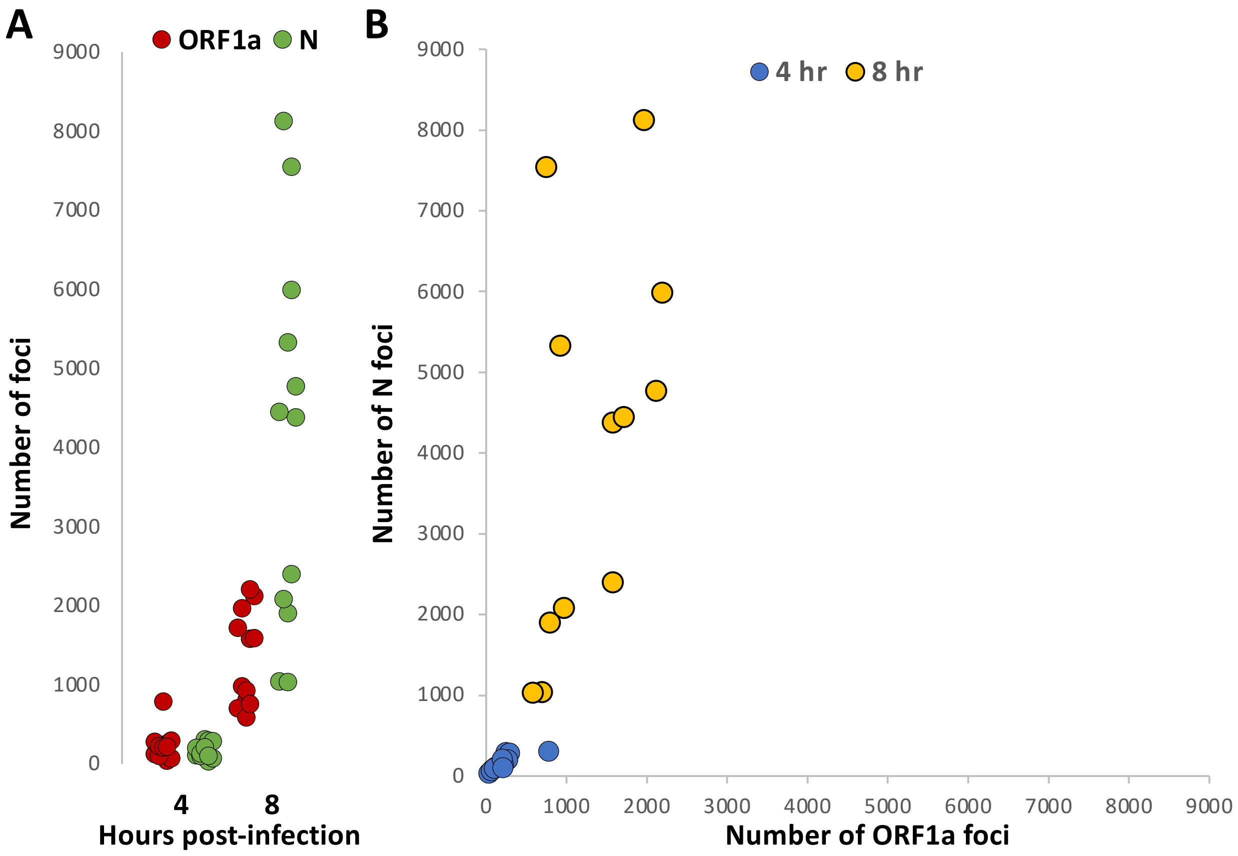


**Figure S2. SARS-CoV-2 genome and mRNA abundance in single-cells at early times post-infection.**
(A) smFISH for ORF1a and N regions of SARS-CoV-2 four- and eight- hours post-infection with SARS-CoV-2. (B) Scatter-plot quantifying smFISH for ORF1a (x-axis) and N region (y-axis), which captures sub-genomic RNAs, at indicated times post-infection. At four hours post-infection, N-targeted RNAs are equivalent to full-length genomes, but are more abundant than full-length genome at eight hours post-infection.

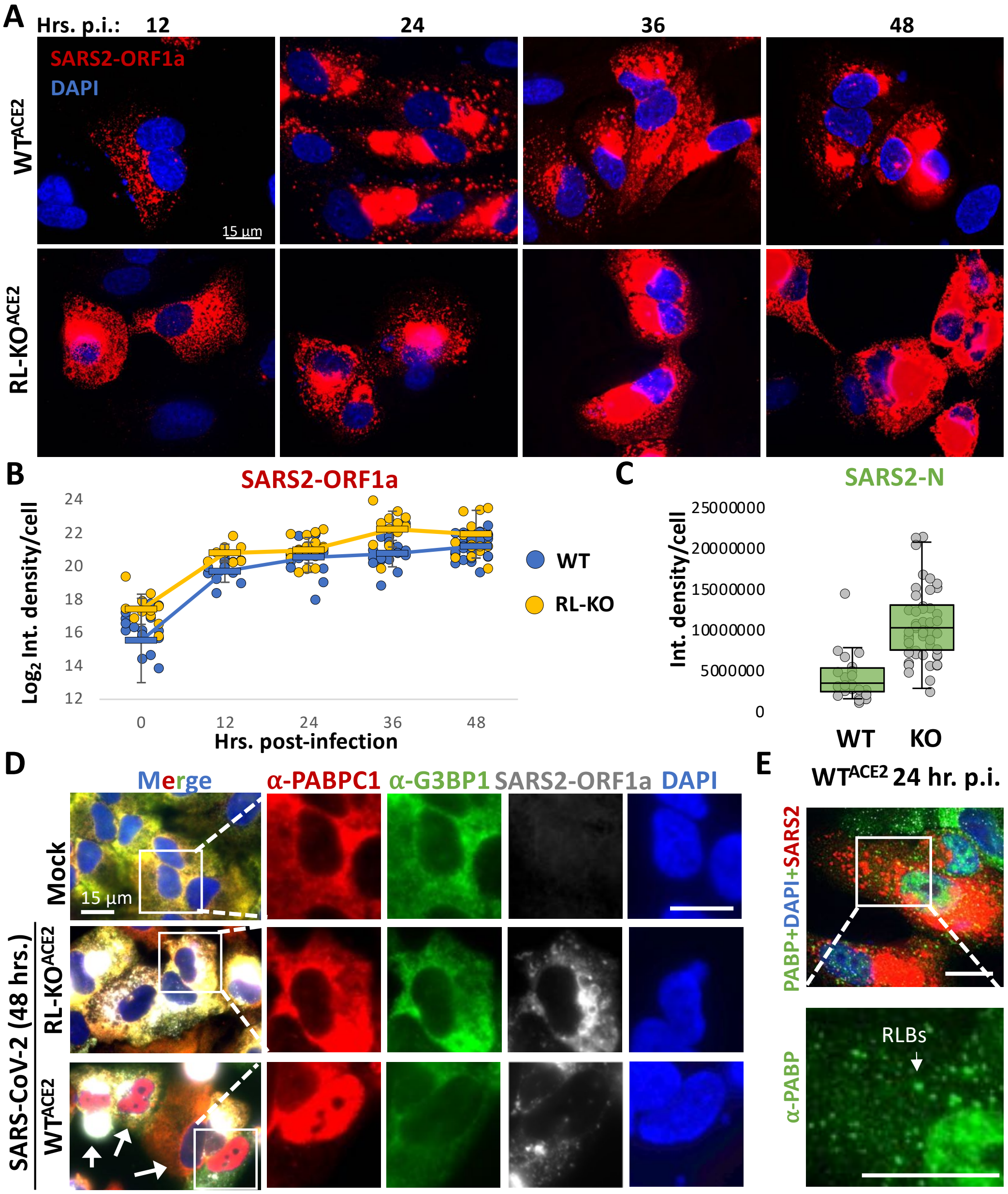


**Figure S3. RNase L is activated by SARS-CoV-2 infection.**

(A) SARS-CoV-2 full-length genome (ORF1a probes) at indicated times post-infection WT$^{ACE2}$ and RL-KO$^{ACE2}$ cells. (B) Quantification of FL genome (ORF1a) fluorescent intensity in WT$^{ACE2}$ and RL-KO$^{ACE2}$ cells. (C) Quantification of fluorescent intensity of sub-genomic RNA (N probes) in WT$^{ACE2}$ and RL-KO$^{ACE2}$ cells. (D) IFA for stress granule markers PABP and G3BP1 and smFISH for SARS-CoV-2 ORF1a in WT$^{ACE2}$ and RL-KO$^{ACE2}$ cells mock-infected or infected with SARS-CoV-2. (E) IFA for PABP and smFISH for SARS-CoV-2 ORF1a in WT$^{ACE2}$ cells.

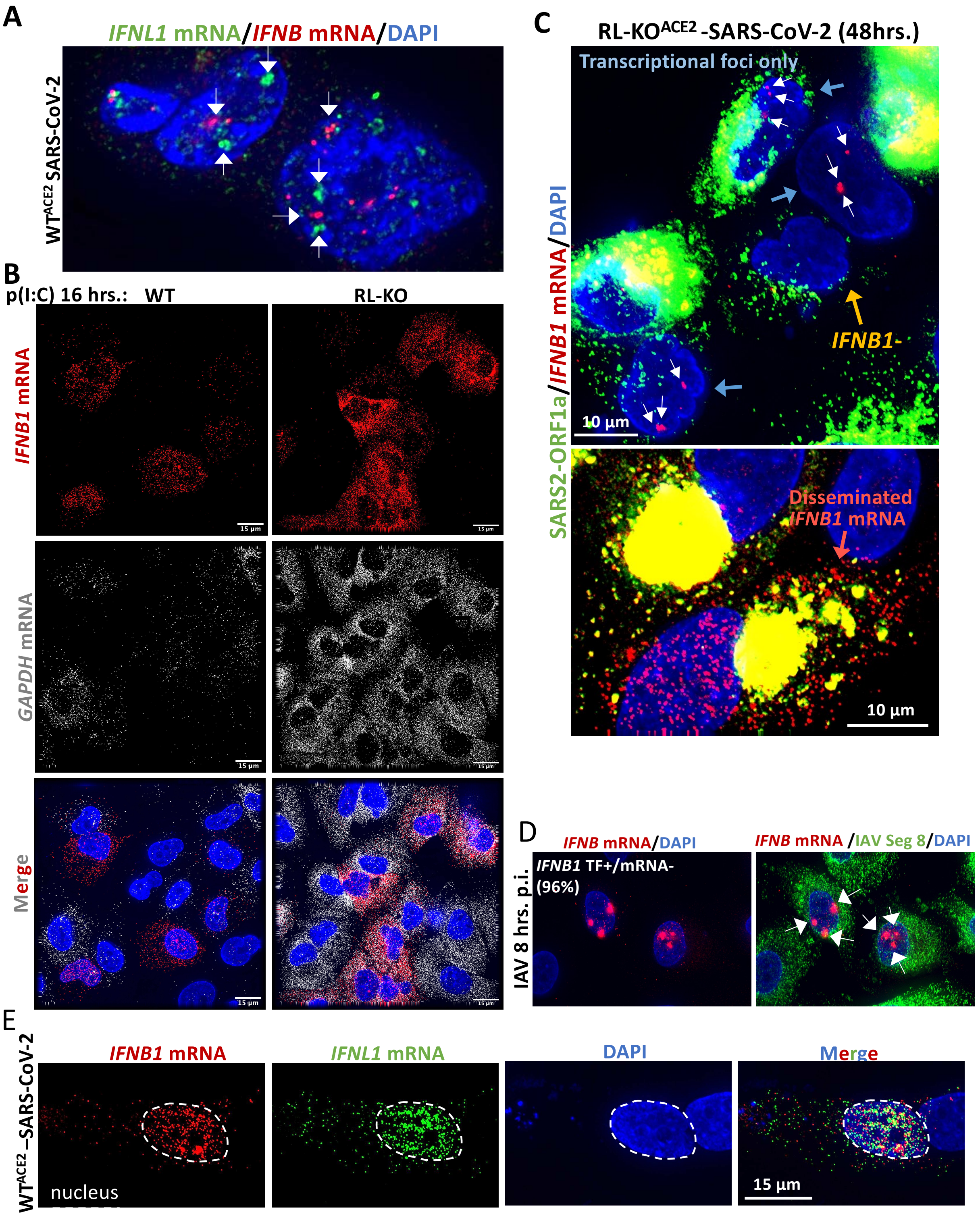


**Figure S4. IFN mRNA localization in SARS-CoV-2-infected and IAV-cells.**
(A) smFISH for *IFNB1* and *IFNL1* mRNAs in WT$^{ACE2}$ cells forty-eight hours post-infection with SARS-CoV-2. (B) Individual smFISH staining for *GAPDH* and *IFNB1* mRNAs shown in Fig. 4B. (C) smFISH for *IFNB1* mRNA and SARS-CoV-2 ORF1a forty-eight hours post-infection in RL-KO$^{ACE2}$. Two fields of view are shown. In the top image, SARS-CoV-2-positive cells stain for *IFNB1* (green arrows), whereas others do not (yellow arrow). Cells that contain

*IFNB1* transcriptional foci (TF) but lack abundant disseminated *IFNB1* mRNA are indicated by blue arrows. The lower image shows a SARS-CoV-2-infected cell that contains abundant and disseminated *IFNB1* mRNA in the nucleus and cytoplasm (red arrow). The yellow staining is due to spectral crossover from the SARS2-ORF1a (green) into the *IFNB1* mRNA (red) channel. (D) smFISH for *IFNB1* and IAV *segment 8/NS* mRNA in WT cells eight hours post-infection with influenza A virus A/Udorn/72 strain (MOI=0.5). Arrows indicate IFNB1 transcriptional foci. (E) smFISH for *IFNB1* and *IFNL1* mRNAs in $WT^{ACE2}$ cells forty-eight hours post-infection with SARS-CoV-2.
